# Nucleus accumbens acetylcholine dips track reward value and temporal costs to shape delay discounting

**DOI:** 10.64898/2026.10.30.760841

**Authors:** Julianna Cavallaro, Dustin R. Zuelke, Czarina M. Maysonet, Ronald E. Salazar, Ruby Setara, Eduardo F. Gallo

## Abstract

As an organism chooses between competing rewards, it must often integrate learned reward value with the changing costs associated with obtaining them. Yet, the brain mechanisms through which these factors jointly guide choice are not well understood. Previous studies have shown that dopamine signals in the nucleus accumbens (NAc) track reward value and are modulated by their associated delay costs. Our prior work further implicated NAc cholinergic interneurons by demonstrating that dopamine D2 receptors in these neurons influence choice under temporal, but not probabilistic, costs. However, whether NAc acetylcholine (ACh) dynamics track reward value and/or delay, and whether they contribute to adaptive choice, remains unknown. Using *in vivo* fiber photometry during a delay discounting task in which mice choose between a small, immediate reward and a larger, delayed reward, we reveal that cue-evoked ACh dips dynamically track relative reward magnitude and are diminished by increasing delays. Our data further show that optogenetically enhancing these dips attenuates the delay-dependent decline in choice of the larger, delayed reward. These findings identify a previously unrecognized role for NAc cue-evoked ACh dips in shaping adaptive choice as temporal costs increase.

## INTRODUCTION

The ability to adaptively select which rewards to pursue as their relative costs change helps organisms direct their behavior toward beneficial outcomes. However, choices may become maladaptive when immediate rewards are disproportionately favored at the expense of future rewards. These behavioral alterations are observed in substance use disorders, attention-deficit hyperactivity disorder, and schizophrenia, compromising daily functioning and long-term clinical outcomes [1,2]. Yet, the fundamental brain mechanisms that integrate reward value and temporal costs to guide adaptive choice remain poorly understood.

Intertemporal choice is typically measured in humans and animals using delay discounting tasks which quantify the degree to which delayed rewards are devalued or “discounted” relative to immediate rewards [3–5]. Lesions and imaging approaches have implicated various brain regions in delay discounting, including the prefrontal cortex, basolateral amygdala, and nucleus accumbens (NAc) [6–8]. Furthermore, behavioral neuropharmacological studies in animals have suggested causal roles for dopamine and acetylcholine (ACh) transmission in delay discounting [9–13]. However, these strategies can only provide limited insight into the specific circuits and dynamic processes through which dopamine and ACh signals influence value-based decision-making [14,15].

Nonetheless, recent work using fast-scan cyclic voltammetry and optogenetics indicated that cue-evoked dopamine release in NAc can alter delay discounting by encoding delay-dependent changes in reward value [16,17]. Our previous work further implicated NAc cholinergic interneurons (CINs) in this process by demonstrating that alterations in dopamine D2 receptor (D2R) expression in these neurons bias choice under temporal, but not probabilistic, costs [18]. However, it remains unknown how ACh dynamics in NAc contribute to value-guided choice as delay costs change.

CIN-derived ACh is known to exert broad influence over local DA release and corticostriatal plasticity [19–28]. In addition, seminal work demonstrated that reward-associated cues trigger multiphasic firing responses in tonically active neurons, widely assumed to be CINs, in the primate dorsal striatum [29,30]. These cue-evoked patterns of CIN activity have been reported across species and striatal subregions and feature a distinct pause in firing that is variably flanked by short firing bursts [29–33]. While these patterns can be evoked by salient or aversive stimuli, the pause triggered by reward-predictive stimuli displays unique features, becomes more prominent with repeated cue-reward associations, and can coincide temporally with cue-evoked dopamine neuron activity [34–36]. As such, the cue-evoked cholinergic pause and the resulting reduction in ACh have long been hypothesized to play permissive roles in dopamine-dependent plasticity and reinforcement learning [36,37]. Because reward-predictive cues elicit information about expected outcomes that is relevant for subsequent choice behavior [38], we hypothesized that cue-evoked ACh “dips” in the NAc contribute to shaping how learned reward value guides choice as delay costs increase.

Here, using *in vivo* fiber photometry to record NAc ACh signals in mice during a delay discounting task, we found that cue-evoked dips in ACh not only track the value of rewards of different magnitude, but are also diminished by progressively longer delays to the larger reward. We further show that optogenetic enhancement of the cue-evoked ACh dip predicting the delayed reward is sufficient to reduce delay discounting. Together, these findings identify cue-evoked NAc ACh dips as a dynamic cholinergic signal that shapes how learned reward value is used to guide choice when facing increased temporal costs.

## METHODS

### Animals

Adult male and female ChAT-Cre mice (B6.FVB(Cg)-Tg (Chat-cre) GM60Gsat/Mmucd, GENSAT; MMRRC stock no. 030869-UCD) [39], were used in fiber photometry experiments and ChAT-IRES-Cre mice (ChAT^tm1(cre)Lowl^/MwarJ; JAX stock #031661)[40] were used in optoinhibition experiments. Both lines were hemizygous for the allele and maintained as C57BL/6J-congenic strains. Mice averaged approximately 6.5 months of age and were age-matched across groups. Mice were housed in groups of 3–5 per cage, whenever possible, on a 12-h light/dark cycle, and all experiments were conducted during the light cycle. All experimental procedures were performed in accordance with NIH guidelines and were approved by the Institutional Animal Care and Use Committees of Fordham University.

### Surgical procedures

Mice underwent stereotaxic surgery under isoflurane-induced anesthesia and received adeno-associated virus (AAV) infusions into the NAc. For fiber photometry, AAV-GRAB_ACh3.0_(ACh4.3) (WZ Biosciences; lot 20201222) and AAV5-Ef1a-DIO-mCherry (UNC Vector Core; lot AV4311G) were mixed 1:1 (total vol. 360 nl) and co-injected using Bregma-based coordinates: AP, + 1.7 mm; ML, ±1.2 mm; and three DV sites, −4.2, −4.1, −4.0 mm (from dura). A fiber optic cannula (Doric, MFC_400/430-0.66_5mm_MF2.5_FLT) was unilaterally implanted at DV = −4.1 mm. For optoinhibition, AAV9-GRAB_ACh3.0_(ACh4.3) was mixed 1:1 and bilaterally co-injected with either AAV5-Ef1a-DIO-eNpHR3.0-mCherry (UNC Vector Core; lot AV4832G) or AAV5-Ef1a-DIO-mCherry at AP, +1.7; ML, +/−1.25; DV −4.2, −4.1, −4.0 mm. Dual optic fibers (Doric, TFC_400/430-0.66_5mm_TS2.5_FLT) were positioned at DV = −4.1. AAV assignment was counterbalanced for sex, age, and home cage origin.

### Behavioral apparatus

Light and sound-attenuating operant cabinets containing a behavioral chamber (Med-Associates, Env-307w) with interior flooring consisting of metal rods spaced 0.87 cm apart and an open ceiling were used in all behavioral experiments. One wall of the chamber contained a central feeder trough flanked on both sides by retractable levers with cue lights above. The central feeder trough was connected to a liquid dipper arm that, when raised, delivers (13-15 μl) of undiluted evaporated milk. Head entries into the trough were recorded with an infrared (IR) photocell detector. A house light located opposite the trough illuminated the chamber during behavioral sessions. The experimental protocols were controlled via Med-PC computer interface and Med-PC V software (Med Associates).

### Delay discounting

Mice were trained on a two-lever delay discounting task, as previously described [18]. Behavioral training began 4 weeks following viral infusions. Mice were food-restricted and maintained at 85–90% of their baseline body weight, with water available *ad libitum*. Mice first were trained to criteria for dipper retrieval and lever pressing at fixed ratio-1 (FR-1) schedule on both levers, as described. Once 30 reinforcers were earned on each lever, initial lever preference was assessed by presenting both levers over 3 sessions. After successful completion of the criteria for trough and lever press training as previously described the lever with the greater total number of presses assigned as the mouse’s initially preferred lever. The preferred lever was subsequently assigned the small reward (1 dipper), while the less preferred lever was assigned the large reward (3 milk dippers) to counteract baseline side biases and establish a learned preference for the large reward during the initial “No-Delay” phase.

In the initial “No-Delay” phase, mice received 14 daily sessions where the delay to delivery of both the small and large milk rewards was set to 0 s following a lever press. Sessions began with 20 forced choice trials where a single lever was presented and rewarded on an FR-1 schedule. A lever light appeared for 5 s before the extension of the associated lever. Mice received 10 “Large” and 10 “Small” forced trials in alternating order. In the remaining 20 “free choice” trials, both lever lights were illuminated for 5 s followed by extension of both levers. Levers remained extended until a press on either lever was made. New trials began following a variable intertrial interval (ITI) (mean = 30.3s). The No-Delay sessions were followed by delay discounting sessions in which the delay to the large reward was increased across sessions, while the delay to the small reward remained at 0 s. Time delays (2, 4, 6, 8, and 10 s) to the large reward following a lever press were presented in separate sessions (3 sessions for each delay) in ascending order [18]. Large reward preference was determined as a percentage of free choice trials in which the large reward lever was chosen on the final session at each delay, unless otherwise noted. Mice that did not achieve ≥50% large choice on the average of the last two sessions of No-Delay training were excluded from the analysis.

### In Vivo Fiber Photometry and Analysis

Fiber photometry (FP) recordings were performed using a TDT LUX RZ-10X processor and Synapse software (Tucker-Davis Technologies). Fiber-coupled LED light at 465 nm and 405 nm was sinusoidally modulated (210/330 Hz), directed to a Doric FMC6 mini-cube, and delivered via low-autofluorescence patch cords to the implanted optical fibers. Light power at the fiber tip was maintained at ∼55 μW (465 nm) and ∼10 μW (405 nm). Emitted fluorescence was collected through the same optical path, demodulated, low-pass filtered at 6 Hz, and sampled at 1017 Hz.

Raw data were processed using GuPPy [41]. To correct for motion artifacts and photobleaching, the 405 nm control signal was fitted to the 465 nm signal using least-squares linear regression and subtracted to generate ΔF/F values, which were smoothed (0.1-s moving average window) and z-scored across the session. Behavioral events from MedPC V were synchronized via TTL pulses. Peri-event time histograms (PETHs) were generated by aligning z-scored signals to cue onset and then baselining each PETH to the 5 seconds before the cue. Biphasic acetylcholine response dynamics were quantified from session-averaged data using peak amplitude and area under the curve (AUC) via the trapezoidal method.

### Optogenetics

Prior to optogenetic testing, mice performed all behavioral training while tethered to a splitter branching fiber-optic patch cord (Doric Lenses; SBP2 _400/430/1100-0.57_FCM-2xZF1.25F). Optogenetic silencing of CIN activity was performed using 593-nm light from a LISER light source equipped with a 593-nm bandpass filter (Doric) controlled by of Doric Neuroscience Studio software. Light was delivered bilaterally into the NAc during the Large cue in forced trials across all delay sessions, starting with the 2-s delay. One second after cue onset, a light pulse (∼0.5 mW) was triggered by a TTL signal from MedPC and sustained for 3 s, followed by a 9-s ramped-down shutoff. The optoinhibition parameters were empirically determined after testing multiple combinations of light power, pulse duration, and ramp-down length in pilot mice co-expressing eNpHR3.0 and ACh3.0 in NAc. We selected the configuration that required the minimum power to significantly suppress the ACh signal for approximately 4 s without inducing a rebound effect. To verify successful reduction of ACh fiber photometry signals in the experimental mice in this study, the 593-nm light source was connected with a bifurcating cable (Doric, MFP_600/630/LWMJ-0.48_FCM-FCM) to 2 separate iFMC6 minicubes, each independently connected to one hemisphere via fiber patch cords (Doric, MFP_400/430/1100-0.57_FC-ZF1.25(F)_LAF). Prior to the start of behavior training, each mouse received a short test (10 x 3-s pulses, 48 s ITI) using the optoinhibition parameters above. In addition, at the conclusion of experiments, we confirmed effective and temporally-restricted optoinhibition during the Large cue in additional delay discounting sessions.

### Immunofluorescence

Mice were transcardially perfused with ice-cold 4% paraformaldehyde (Fisher Scientific) in PBS under deep anesthesia (100 mg/kg ketamine and 10 mg/kg xylazine, i.p.). Fixed brains were harvested, postfixed overnight, and washed in PBS before 50-μm coronal sections were prepared with a vibratome (Leica, VT1200S). For immunohistochemistry sections were washed 3x in TBS and then incubated for 2 h at room temperature in blocking solution (10% fetal bovine serum, 0.5% Triton X, 0.5% bovine serum albumin in TBS). Sections were labeled overnight at 4°C with primary antibodies against dsRed to label mCherry (Takara, rabbit anti-dsRed, 1:500; 632469) and GFP to label GCaMP (Abcam, chicken; 1:1000; AB13970). Sections were incubated with corresponding fluorescent secondary antibodies (Thermo Fisher Scientific, donkey anti-chicken Alexa 488; 1:400; A78948 and donkey anti-rabbit Alexa 546; 1:400; A10040) for 2 h at room temperature, washed, mounted on slides, and coverslipped with Vectashield Hardset with DAPI (Vector Laboratories, H-1200-10). Images were obtained with a Nikon Eclipse Ti-E epifluorescence microscope and processed with NIH Image J software. The position of implanted fibers was estimated by manual alignment to the Franklin and Paxinos brain atlas (3rd edition).

### Data analysis

Sample sizes were determined by performing statistical power analyses based on effect sizes observed in our prior work or similar work in the literature. Statistical analyses were performed using Prism 11 (GraphPad) or SPSS. For all analyses, data are expressed as the mean ± standard error of the mean (SEM), unless noted otherwise noted and p values < 0.05 were considered statistically significant. Analyses of two-group data across delays or sessions were done using two-way repeated measures ANOVA or mixed-effects models, followed by *post hoc* comparisons of within-group and cross-group differences using Tukey’s HSD test. One-way repeated measures ANOVA was used to assess overall differences across training sessions or experimental conditions. We used using Prism’s test for linear trend to evaluate progressive changes across ordered sessions or delays. Pairwise comparisons were performed using t-tests for normally distributed data or Wilcoxon signed-rank or Mann-Whitney U tests for data that did not meet normality assumptions. Pearson’s *r* correlation was used to evaluate the relationship between percent large preference and Large cue or Small cue dips. A linear mixed-effects model was used to test whether reward cue type was associated with cue-evoked ACh AUC after accounting for press latency. Reward cue type and press latency were included as fixed effects, with subject included as a random intercept to account for repeated trials within each animal. Significance of fixed effects was assessed using Type III tests. Investigators were blinded to experimental conditions throughout behavioral assays and data analysis.

## RESULTS

### Cue-evoked ACh dips emerge with experience and track reward size

To begin to unravel how NAc CINs contribute to delay-based decision-making, we first sought to characterize NAc ACh dynamics using *in vivo* fiber photometry during a delay discounting task adapted for mice [18,42,43]. To this end, we expressed the genetically-encoded GPCR-Activation Based ACh sensor (GRAB_ACh3.0_)[44] in NAc of ChAT-IRES-Cre mice, together with a Cre-dependent mCherry construct for *post hoc* histological visualization of CINs **(Fig. 1A)**. Fluorescent NAc ACh signals were recorded unilaterally via an optic fiber cannula throughout behavioral sessions (**Fig. S1**).

**Figure 1.**
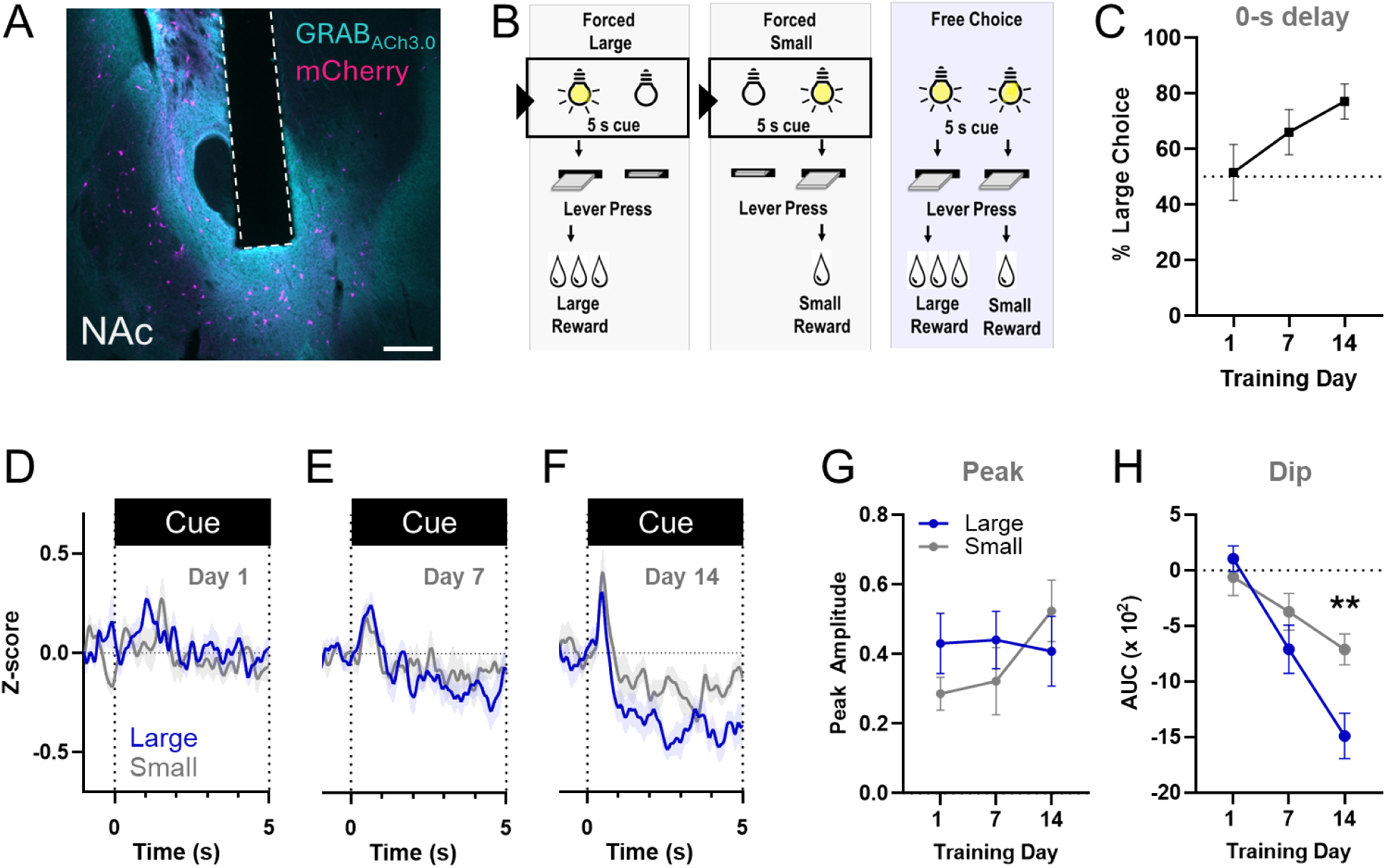
Cue-evoked ACh dips emerge with experience and track reward size. **A** Representative co-immunolabeling showing GRAB_ACh3.0_ sensor expression and mCherry-positive CINs in NAc of ChAT-Cre mice. Dotted line depicts optic fiber tract. Scale = 0.4 mm. **B** Overview of the No-Delay training session structure. Each session was comprised of 20 forced followed by 20 free choice trials. All trials began with a 5-s cue light above the lever(s) prior to lever extension. In free choice trials, both cue lights were simultaneously presented, followed by extension of both levers. Fiber photometry ACh signals were aligned to the onset of the 5-s cues in forced trials, as depicted by the bounding boxes. Illustration adapted from Saddoris et al. *Biol Psych*. 2015 [17]. **C** Large-reward choices across training days. While a one-way RM ANOVA did not reach statistical significance (main effect F _(2,18)_ = 2.964, p = 0.077), there was a significant linear trend (F _(1,18)_ = 5.890, p = 0.026), demonstrating a progressive increase in preference for the large reward across training. **D-F** Mean ± SEM GRAB-ACh z-scored signals aligned to cue onset during Small and Large forced trials measured during No-Delay training. **G** Two-way RM ANOVA revealed no significant effect of cue type or training in ACh peak amplitude measured in the 1^st^ second from cue onset. Delay: F _(1.722, 30.99)_ = 1.246, p=0.2970; reward type: F _(1, 18)_ = 0.3127, p=0.5829; delay x reward type: F _(1.722, 30.99)_ = 1.989, p=0.1589. **H** Mean cue-evoked ACh dip (negative AUC 1-5 s from cue onset) significantly increased across No-Delay training: F _(1.990, 35.82)_ = 26.10, p<0.0001, with a significant training x cue type interaction (F _(1.990, 35.82)_ = 4.614, p=0.0166). Tukey’s post hoc: Day 14 Large vs Small-reward cues: **p = 0.0063. For all panels, N = 10 mice.

Mice were trained to choose between pressing one of two levers that would deliver either a small reward or a 3X larger reward without delays. Each session began with 20 forced choice trials (10 ‘Small’, 10 ‘Large’), where a 5-s cue light preceded the extension of a single active lever that delivered its assigned reward. This was followed by 20 free choice trials, where both cue lights were simultaneously presented for 5 s prior to the extension of both levers, allowing mice to choose either reward. **(Fig. 1B)**. After 14 days of this ‘No-Delay’ training phase, the mean Large-reward choice increased from 51.5 % on day 1 to 77.0 % on day 14, as previously observed [18] **(Fig. 1C)**.

Predictive cues influence choice behavior [45,46] and elicit distinct multiphasic patterns of striatal CIN firing, including bursts and pauses [30]. Therefore, we first determined whether cue-evoked responses in NAc ACh would develop as mice learned to associate cues with the Large or Small-reward options in the No-Delay phase. To this end, NAc ACh3.0 photometry signals were aligned to the onset of each cue type in forced trials on days 1, 7, and 14. Onset of either cue elicited a transient increase above baseline, followed by the development of a sustained dip in ACh below baseline that persisted through the end of the 5-s cue period **(Fig. 1D-F)**. While the initial peak response was not significantly altered by training or by cue type **(Fig. 1G)**, there was a significant increase in the ACh dip magnitude **(Fig. 1H)**. Moreover, the Large cue elicited a significantly greater ACh dip than the Small cue by day 14, matching the behavioral preference for the large reward on this day. Because Large and Small-reward trials differed in subsequent press latency **(Fig. S2),** we tested whether this difference accounted for the greater cue-evoked ACh response associated with Large cues. The reward type effect remained significant after controlling for press latency in a linear mixed-effects model (F_(1,197)_ = 13.976, p < 0.001). Together, these results suggest that cue-evoked ACh dips emerge as animals learn the significance of reward-predictive cues, and that the magnitude of the learned ACh dip response tracks the size of the predicted reward.

### Delay costs attenuate ACh dips evoked by Large-reward cues

We next determined whether increasing delays to the large reward would diminish cue-evoked ACh responses. In the delay discounting sessions, which had the same structure as the preceding No-Delay training, the large reward was delivered after a delay that increased across sessions [18] **(Fig. 2A)**. We analyzed ACh signals evoked by Large and Small cues in forced trials during the last session at each delay. As anticipated, the percent Large choice significantly decreased as the delay to the large reward increased **(Fig. 2B)**. **Figure 2C** shows mean ACh responses to either cue as a function of delay to the large reward. The initial cue-evoked ACh peak showed no significant effects of cue type or delay **(Fig. 2D)**. However, the contrast between Large and Small ACh dips that was evident in the 0-s and 2-s delay sessions was reduced at longer delays, primarily due to attenuation of the Large-cue dip **(Fig. 2E)**. Moreover, the dynamic ACh dip response mirrors the behavioral shift towards the small, immediate reward and was similarly observed in both males and females **(Fig. S3).** These findings suggest that the ACh dips evoked by the Large cue are sensitive to temporal costs associated with reward delivery.

**Figure 2.**
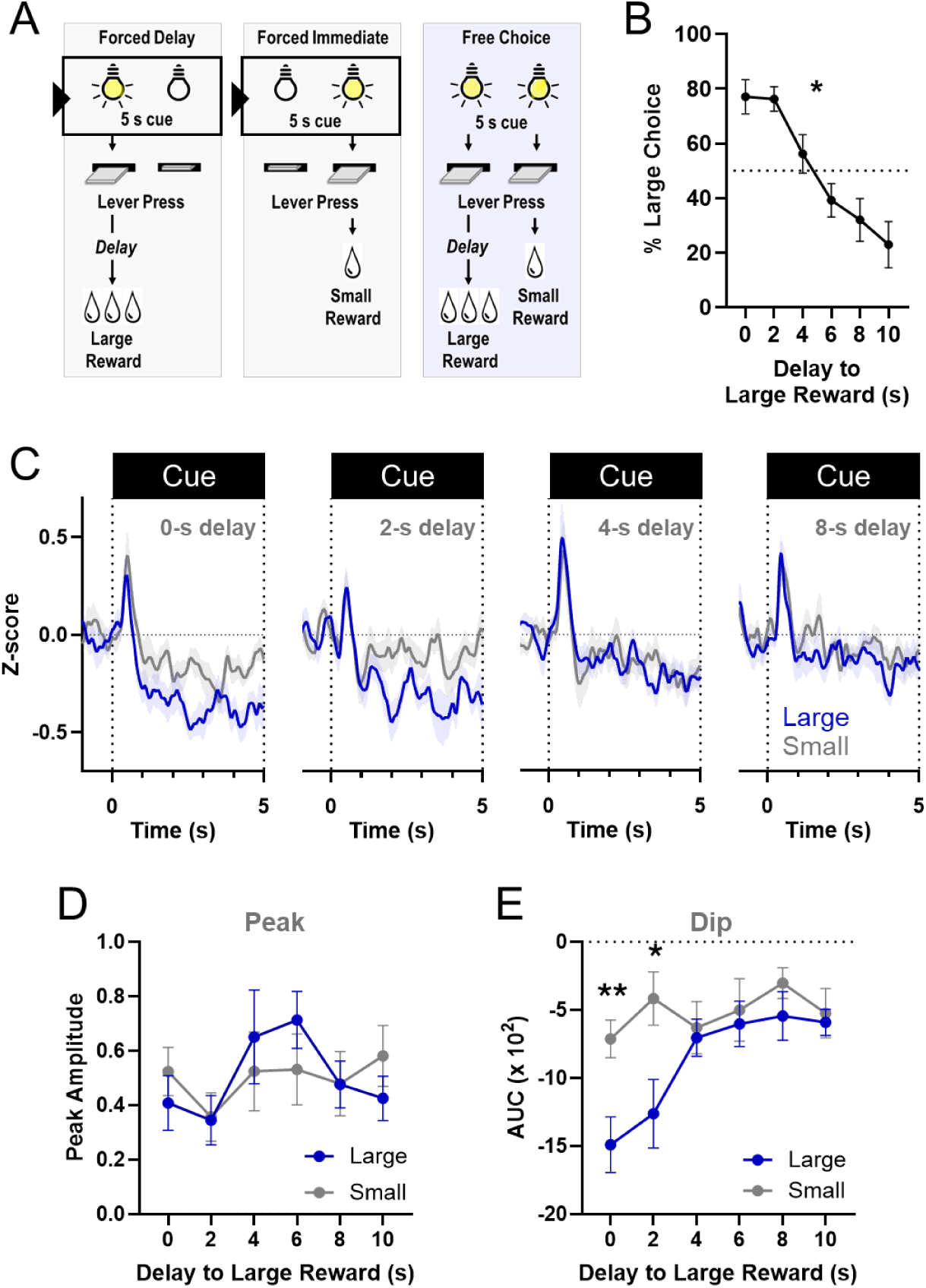
Delay costs attenuate ACh dips evoked by Large-reward cues. **A** Structure of the delay discounting session, which consisted of 20 forced trials followed by 20 free choice trials. Pressing the Large lever resulted in large rewards delivered after a delay (see Methods). ACh signals were aligned to the onset of Large or Small cues in forced trials (bounding boxes) **B** One-way RM ANOVA showed a significant effect of delay on % Large choices (F_(5, 45)_ = 25.66, *p < 0.0001). **C** Mean GRAB-ACh z-scores aligned to forced cue onset showed cue-evoked ACh peaks and dips during delay discounting sessions. **D** No significant effect of delay or cue type was detected for the initial ACh peak amplitude following forced cue presentation. Two-way RM ANOVA, Delay: F _(3.492, 62.86)_ = 2.426, p = 0.0650; Cue type: F _(1, 18)_ = 0.001094, p=0.9740; Delay x Cue type: F _(3.492, 62.86)_ = 1.134, p=0.3459. **E** The mean ACh dip AUC evoked by Large and Small forced cues differed significantly depending on the delay to the large reward. Two-way RM ANOVA: significant main effects of delay (F _(4.085, 73.52)_ = 5.142, p=0.0010) and cue type (F _(1, 18)_ = 4.864, p=0.0407), and a significant delay x cue type interaction (F _(4.085, 73.52)_ = 2.826, p=0.0298). Tukey’s post hoc comparisons showed significant differences between Large and Small cues at the 0-s delay and 2-s delay (**p = 0.0063; *p = 0.0169). For all panels, N = 10 mice.

### Large cue-evoked dips in forced trials correlate with subsequent Large-reward preference

We next sought to determine whether cue-evoked ACh neural responses observed in the forced trials are related to subsequent choice behavior in the delay discounting task. We performed a Pearson correlation analysis to examine the relationship between the magnitude (AUC) of the ACh dip evoked by either the Large or Small cues in forced trials and the percent preference for the large reward for each animal at each delay. We found a significant positive relationship between Large-cue dip magnitude and the percent Large-reward preference **(Fig. 3A).** In contrast, no significant correlation was found for the Small cue **(Fig. 3B)**. These findings suggest that the magnitude of the Large cue-evoked ACh dip in forced trials is related to how strongly animals continue to prefer the large reward as its delivery is delayed.

**Figure 3.**
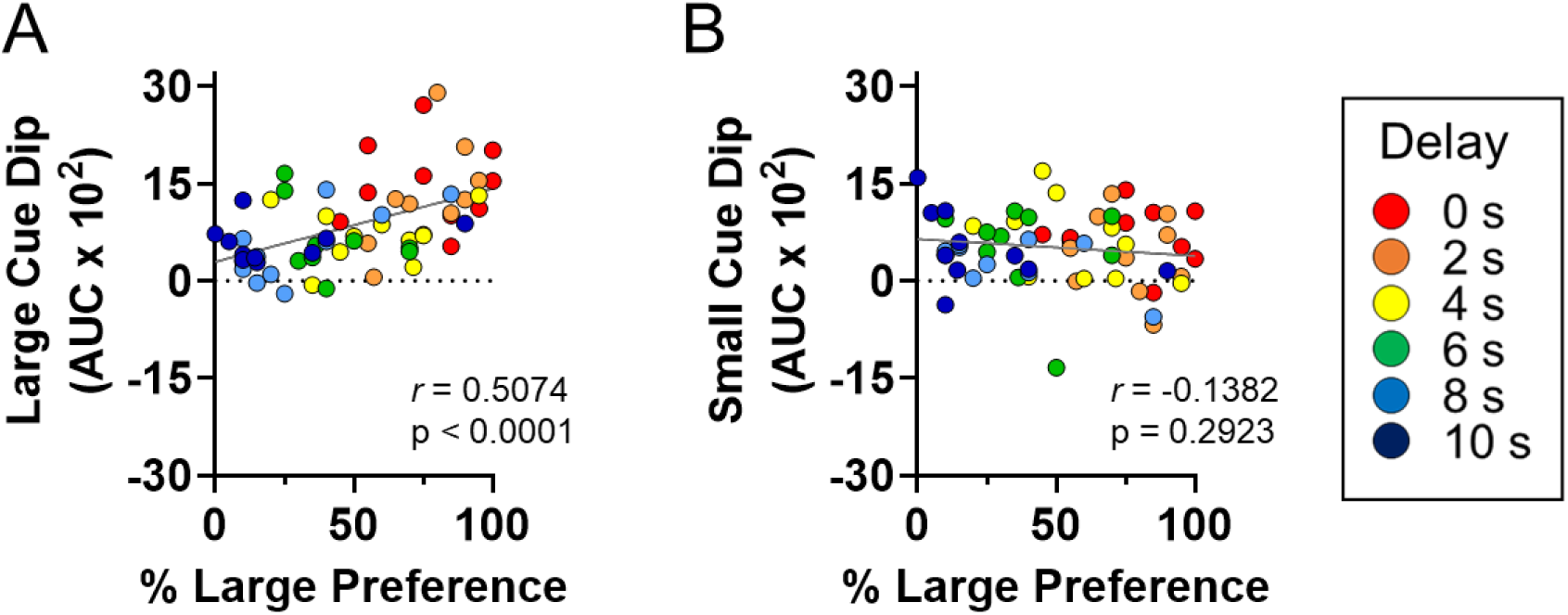
Large cue-evoked dips in forced trials correlate with subsequent Large preference. **A** Scatter plot showing a significant correlation between the magnitude of the ACh dip evoked by the forced Large cues (absolute value AUC) and the behavioral preference for the large reward subsequently expressed in the choice trials for each animal across delays. **B** In contrast, there was no significant relationship between small cue dip and percent large reward preference. N = 10 mice.

### Cue-evoked ACh dips in free choice develop with training and do not distinguish between subsequent choices

In forced trials, the cue predicted a specific reward and its corresponding action without requiring a choice between alternative options. We therefore tested whether the cue-evoked ACh signal is similarly expressed in free choice trials, where animals encounter both cues simultaneously and can select either the large or the small reward. First, we analyzed ACh responses during the 14-day No-Delay phase **(Fig. 4A-D)**. Cue-evoked ACh dips in free choice trials were nearly absent on day 1, but significantly increased in magnitude by days 7 and 14 **(Fig. 4B-C and S3E-F)**, reminiscent of the ACh dip responses observed for the Large cue in the forced trials **(Fig. 1H).** The initial positive peak also showed a significant increase with training **(Fig. S4A)**. We further sorted cue-evoked ACh trial data based on the eventual choice for large or small rewards. A mixed-effect analysis revealed that both reward options were associated with a training-dependent increase in cue-evoked ACh dip magnitude **(Fig. 4D)**. However, there was no significant effect of chosen reward or a significant training day-by-chosen reward interaction.

**Figure 4.**
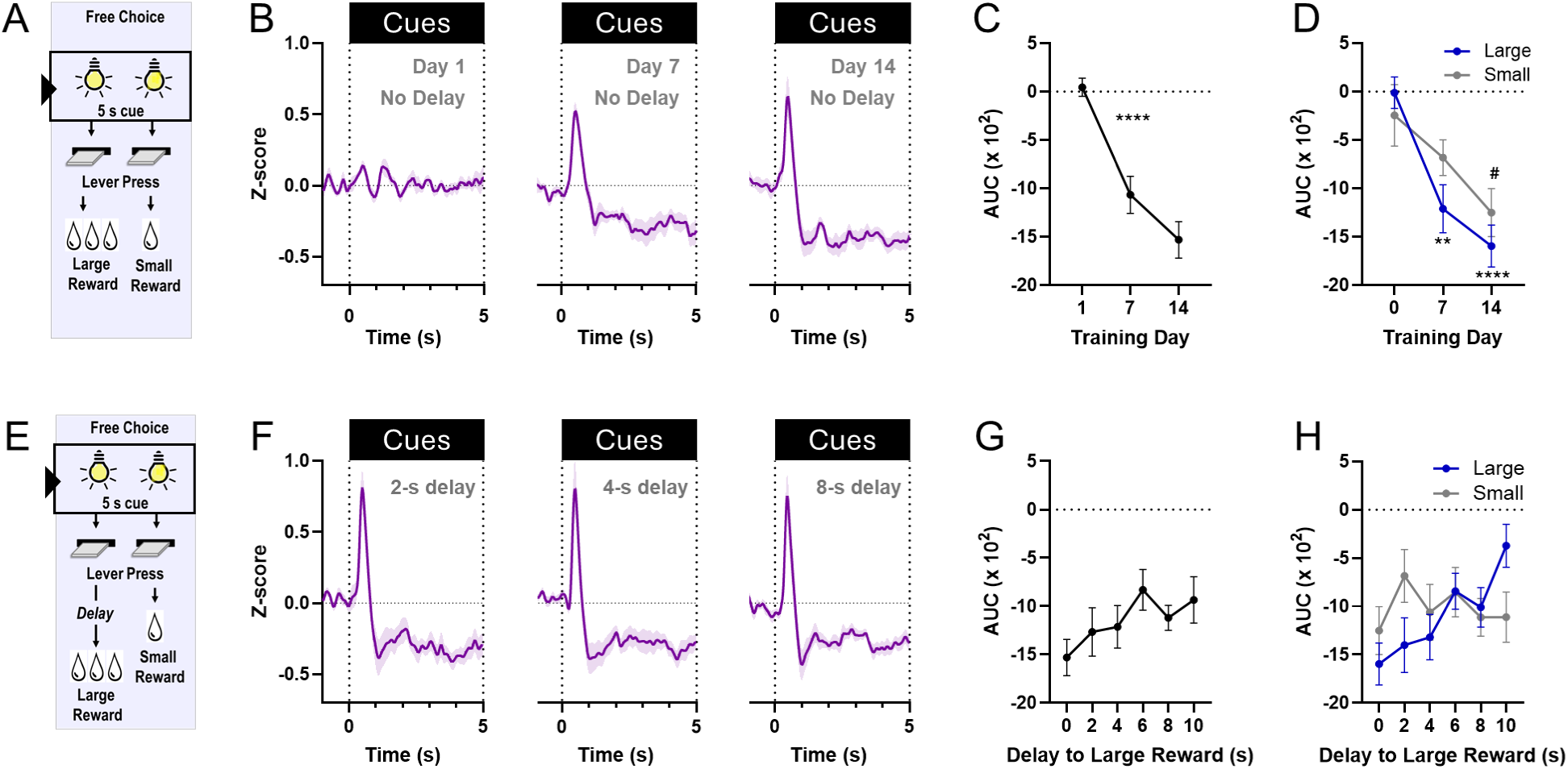
Cue-evoked ACh dips in free choice develop with training and do not distinguish between subsequent choices. **A** In free choice trials during No-Delay training, both cues were simultaneously lit for 5 s, followed by extension of the levers. After the lever choice, rewards were delivered immediately. ACh signals were aligned to dual cue onset. **B** Mean GRAB-ACh z-scored signals aligned to free cue onset across No-Delay training. AUC was analyzed from 1-5s following cue onset. **C** ACh dips following dual cue onset significantly increased with No-Delay training (one-way RM ANOVA, F _(2, 18)_ = 28.61, ****p<0.0001). **D** Cue-evoked ACh dip data from free choice trials was sorted by subsequent reward choice. Mixed effects model: delay effect (F _(2, 33)_ = 16.30, p < 0.0001), but no effect of the reward choice (F _(1, 18)_ = 1.207, p = 0.2863) or training day x reward choice interaction (F _(2, 33)_ = 1.521, p = 0.2334). Tukey’s post hoc: Large choice: day 1 vs. 7 (**p = 0.0016), day 1 vs. 14 (****p < 0.0001). Small choice: the preceding cue-evoked dip was larger at Day 14 vs. Day 1 (^#^p = 0.0134). **E** In the delay discounting phase, both light cues were illuminated for 5s. Pressing the Large lever delivered the large reward after a delay that increased across sessions (0-10 s); pressing the other lever delivered a small, immediate reward. **F** ACh signals aligned to dual cue onset on the last day of 2, 4, and 8-s delay sessions. **G** ACh dips evoked by dual cue presentation were not significantly altered by delay overall (one-way RM ANOVA, F _(5,45)_ = 1.981, p = 0.099), but there was a significant linear trend: F _(1,45)_ = 6.6549, p = 0.0139. **H** Mixed effects analysis examining the effects of delay and eventual choice (Large vs. Small) revealed no significant main effects of delay (F _(3.402, 59.87)_ = 2.363, p = 0.0727); or choice (F _(1, 18)_ = 0.1906, p = 0.6676), and a trend toward a delay × choice interaction (F _(3.402, 59.87)_ = 2.553, p = 0.0569). For all panels, N = 10 mice.

We then analyzed cue-evoked ACh signals in the free choice trials of the delay discounting phase, where the delay to the large reward increased across sessions **(Fig. 4E).** ACh dips evoked by simultaneous cue presentation were pronounced and showed a significant linear trend with increasing delays to the Large reward, although the overall effect of delay did not reach significance **(Fig. 4F-G)**. The initial positive peak was not altered by delays **(Fig. S4B)**. After sorting cue-evoked trial data for both free cues by the reward ultimately chosen, a mixed effects model found no significant main effect of reward choice or delay on ACh dip magnitude **(Fig 4H).** While the interaction between reward choice and delay approached statistical significance (F_(3.402, 59.87)_ = 2.553, p = 0.0569), with AUC values trending toward a decrease in dips preceding Large choices, our data did not provide clear evidence of this distinction **(Fig. 4H)**. Together, these findings suggest that dual reward cues in free choice trials evoke robust, experience-dependent ACh dips that do not clearly distinguish subsequent Large from Small choices across delays.

### Enhancing the Large cue-evoked ACh dip maintains behavioral preference for the large, delayed reward

Our data thus far indicate that the Large-reward cue evoked a larger ACh dip than the Small-reward cue at 0-s and 2-s delays, when the behavioral preference for the large reward was high, but was progressively attenuated at longer delays as the large reward preference declines. Therefore, we hypothesized that if the Large cue-evoked ACh dip plays a role in maintaining the value representation of the large reward across delays, then enlarging the cue-evoked dips for the Large option should prevent their delay-dependent reduction and decrease discounting.

To this end, we co-infused AAVs expressing DIO-eNpHR3.0-mCherry (or DIO-mCherry) and GRAB_ACh3.0_, followed by bilateral optic fiber implantation into the NAc of ChAT-Cre mice **(Fig. 5A and Fig. S5A)**. To specifically augment the cue-evoked ACh dip as animals experienced increasing delay costs, we optogenetically inhibited NAc CINs starting 1 s after Large cue onset in forced trials across all delay sessions **(Fig. 5B)**. We confirmed that our protocol selectively reduced NAc ACh without inducing a rebound effect and enlarged the cue-evoked ACh dip in Large forced trials in eNpHR3.0 mice **(Fig. S5B-D)**. In addition, this manipulation did not alter the initial ACh positive peak nor the ACh dip in response to cues in the forced Small or in free choice trials **(Fig. S5C)**. We found that restricting 593-nm light illumination to forced Large cues led to significantly reduced delay discounting in eNpHR3.0 mice compared to mCherry controls, as reflected by their higher average preference for the large reward at longer delays **(Fig. 5C)**. Consistent with this effect, linear discounting slopes were significantly shallower in eNpHR3.0 mice **(Fig. 5C, *inset*)**. These results suggest that enhancing the Large cue-evoked ACh dip is sufficient to reduce the delay-dependent bias toward the smaller, immediate reward.

**Figure 5.**
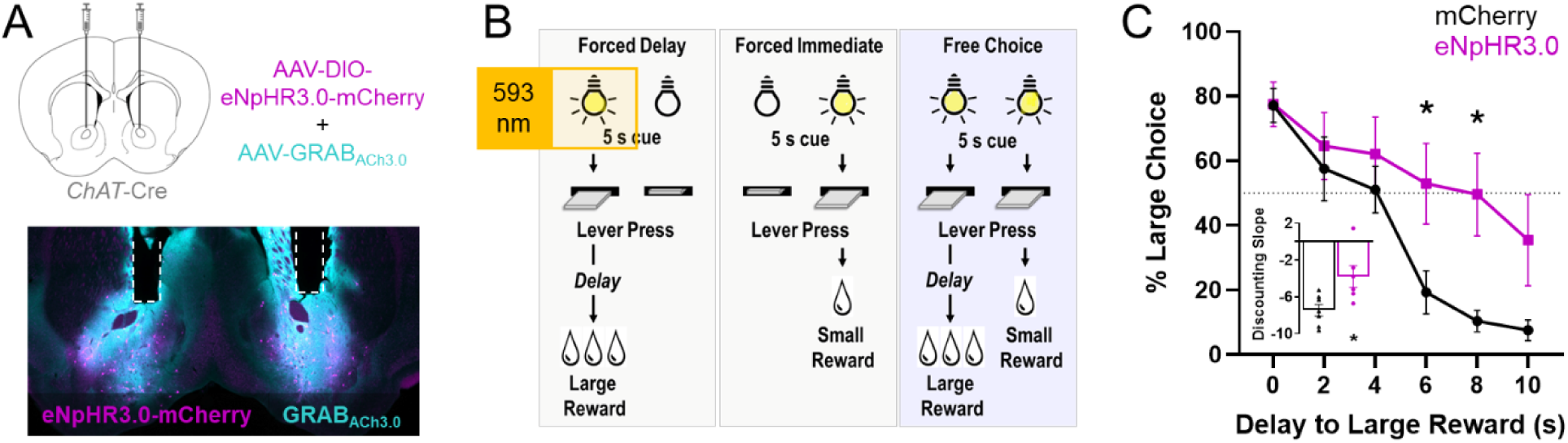
Enhancing the Large cue-evoked ACh dip maintains behavioral preference for the large, delayed reward. **A** *Upper*, mice were bilaterally injected with AAV-GRAB-ACh3.0 and AAV-DIO-eNpHR3.0-mCherry or AAV-DIO-mCherry in the NAc followed by dual fiber-optic cannula implantation. *Lower*, Representative image showing optic fiber locations and co-ACh3.0 and eNpHR3.0-mCherry expression in NAc. **B** Schematic of the delay discounting task with CIN optoinhibition. 593-nm light was applied during Large cues in forced trials, starting 1 s after the cue onset and held constant for 3 seconds and ramped down. **C** CIN inhibition significantly decreased delay discounting in eNpHR3.0 mice compared to mCherry controls. Percent Large preference expressed as an average of the last 2 sessions at each delay. Two-way RM ANOVA: main effect of delay (F _(2.834, 31.18)_ = 34.84, p<0.0001) and a significant delay x virus interaction (F _(2.834, 31.18)_ = 4.804, p=0.0082). Tukey’s post hoc: eNpHR3.0 vs. mCherry at 6-s (*p = 0.0457) and 8-s delays (*p = 0.0264). *Inset*, slopes of best-fit lines relating each mouse’s Large reward preference to delay. Mann-Whitney U test: eNpHR3.0 vs. mCherry (U = 4, *p = 0.014). n = 7 mCherry; 6 eNpHR3.0 mice.

## DISCUSSION

This study, to our knowledge, provides the first characterization of cue-evoked NAc ACh dynamics during delay discounting. We found that cues predicting different reward options evoke sustained ACh dips. These dips emerge through experience and signal the relative value of those rewards. The magnitude of the cue-evoked ACh dip associated with the larger reward was progressively diminished by increasing delays, reflecting the reduced behavioral preference for that reward. Notably, enlarging the ACh dip during cues predicting the large delayed reward attenuated delay discounting. Together, our findings suggest that cue-evoked NAc ACh dips contribute to adaptive choice by influencing how learned reward value guides choice as temporal costs change.

Pioneering *in vivo* electrophysiological recordings showed that repeated cue-reward pairings increase the number of putative CINs that respond with a cue-evoked pause in firing [30]. Subsequent work in the primate dorsal striatum suggested that pauses are most pronounced in response to cues associated with reward, further supporting the notion that they are not solely responses to sensory events [34,35]. In fact, repeated association of an aversive stimulus with subsequent reward can shape the pause response to become more “reward-like,” suggesting that CIN pauses can acquire motivational significance with learning [35]. Our fiber photometry findings extend this framework to ACh dynamics in the mouse NAc. The cue-evoked ACh dip was not clearly present on the first day of No-Delay training, but progressively grew in response to both the Small and Large cues following associative learning. These results are consistent with the emergence of CIN pausing seen in electrophysiology studies [30,35] and with learning-dependent changes in ACh dips in different striatal subregions [47–50].

With increased task experience, mice also showed a greater ACh dip in response to the Large cue compared to the Small cue when large rewards were delivered immediately or after a short delay (2s). This experience-dependent response aligned with the strong behavioral preference for the large reward in choice trials at those delays. Notably, these effects were selective to the cue-evoked dips, as the initial peaks were not significantly affected by training and did not discriminate between reward options. These cue-evoked peaks are likely driven by excitatory cortical or thalamic input, possibly reflecting a sensory response to cue onset [51,52]. These findings suggest that NAc ACh dips are not simply a consequence of cue presentation, but are sculpted through associative learning and reflect differences in the learned value of the rewards predicted by those cues.

We hypothesized that if cue-evoked ACh dips reflect the learned value of available reward options, increasing delays to the large reward delivery would reduce the ACh dip evoked by the Large cue. Indeed, sessions with delays of 4 s or longer displayed a diminished Large cue dip. This is consistent with a weakening of the learned Large value representation, as these delays were also associated with increased behavioral bias towards smaller, immediate rewards. Furthermore, the longer delays made the Large dip comparable to that elicited by Small cues but did not eliminate the dip altogether. This suggests that longer delays to the large reward diminish the ability of the ACh dips to reflect the relative value of the two reward options, while sparing a residual cue-evoked dip associated with the large reward. Moreover, longer delays to the large reward did not translate to enhanced Small cue dips, even though under those conditions the animals increasingly chose the small reward. By primarily tracking the discounting of the large, delayed reward, this pattern suggests that, in forced trials, ACh dips encode temporal cost information in a reward-dependent manner.

Once relative reward value is learned, how does the cue-evoked ACh dip contribute to choice as temporal cost increases? Our correlation analysis implies that information from ACh dips in the forced trials is relevant to behavioral preference in free choice trials (**Fig. 3**). We found that optogenetically enhancing the Large cue-evoked ACh dips in forced trials during delay sessions was sufficient to reduce discounting of the Large reward in free choice trials. The significant delay-by-virus interaction (**Fig. 5**) indicates that the behavioral consequence of enhancing the ACh dip depends on the temporal cost. Indeed, the effect of optoinhibition on choice was readily apparent at delays longer than 4 s, indicating that the dip’s influence is most consequential as temporal costs increasingly diminish the value of the Large reward. These findings suggest that ACh dips shape adaptive choice by influencing how learned value representations are used as temporal costs change.

Our experiments also address the question of how cue-evoked ACh signals contribute to choice when competing cues are presented together. During forced trials, each cue predicts a specific reward and elicits learned information about that reward’s value and associated delay. During free choice trials, simultaneous cue presentation elicits information about both upcoming options. Unlike the differential ACh responses elicited by the separately-presented Large and Small cues, the dual cue-evoked ACh dip in free choice trials did not distinguish the reward that was subsequently chosen. Because the dual-cue ACh dip was robust and qualitatively similar to that evoked by the Large cue alone, it is thus possible that the free choice ACh dip instead reflects the integrated motivational significance of the available reward options. Similar patterns have been reported for cue-evoked dopamine in signaling the best available option in a choice context [53,54]. Our results suggest that ACh dips in response to individual cues do not directly reflect choice, but instead contribute to the learned value representations that are subsequently used to guide choice.

Our results reveal a cue-evoked cholinergic signal within the NAc that is not only sensitive to learned reward value and temporal costs, but that can also causally influence delay-based choice. These new findings align well with reported roles for mesolimbic cue-evoked dopamine in encoding temporally-discounted subjective value [16,17,53–55] and driving choice in similar delay contexts [17]. Together with our recent finding that CIN D2R levels can alter delay discounting [18], these results support the possibility that dopaminergic and cholinergic signaling may dynamically interact within the NAc during cues to shape how learned valued is represented and utilized as temporal costs change. This perspective fits within the hypothetical framework in which increased DA release temporally coincides with ACh dips to modulate corticostriatal plasticity, reinforcement learning, and action selection [36,37,56,57]. Yet, the extent, timing, and functional significance of this NAc DA-ACh relationship in delay-based choice remain to be further investigated.

Our ability to monitor and alter ACh with temporal precision during cues enabled us to isolate an important behavioral contribution of the elusive ACh dip. Similar targeted approaches in awake-behaving animals may uncover other critical information hidden within the complex multiphasic cholinergic patterns observed in response to different behavioral events or contexts [33,47,49,58]. In this regard, a recent study using optogenetic stimulation of rat dorsal striatal CINs to reduce ACh dips during reward delivery found impaired reversal learning [59]. Such temporally-precise strategies, informed by endogenous neural dynamics, may also prove critical for defining functional interactions between different ACh response types and non-cholinergic signals in NAc that support reward-related behavior.

In summary, the present results establish NAc ACh signaling as a key component of the circuit mechanisms underlying delay discounting. By demonstrating that cue-evoked ACh dips can causally influence choice involving delayed rewards, this work identifies a discrete neural signal through which adaptive decision-making can be modulated. Therefore, these findings could be of broad relevance to disorders characterized by maladaptive discounting of future rewards.

## ACKNOWLEDGMENTS

We thank William Mannheim, Alexander Gannon and Gabriela Jammar for technical assistance and Venus Sherathiya for support with Guppy software for fiber photometry data analysis

## AUTHOR CONTRIBUTIONS

J.C, D.R.Z, C.M.M., R.E.S., R.S., and E.F.G. conducted experiments and data analysis. E.F.G. and D.R.Z. wrote the manuscript. E.F.G. designed the experiments. E.F.G. supervised the experiments and data analysis.

