## Supplementary Figures for "Nucleus accumbens acetylcholine dips track reward value and temporal costs to shape delay discounting"

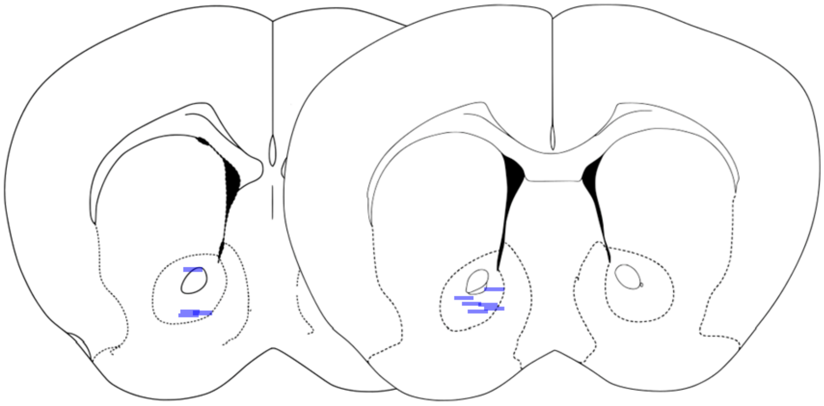


**Figure S1. Fiber optic cannula locations for fiber photometry recordings.** Blue lines represent the approximate locations of individual fiber tips based on the ventral-most point of the fiber track observed across serial histological sections. Numbers indicate distance (mm) from Bregma, according to Paxinos and Franklin mouse brain atlas (3rd ed.).

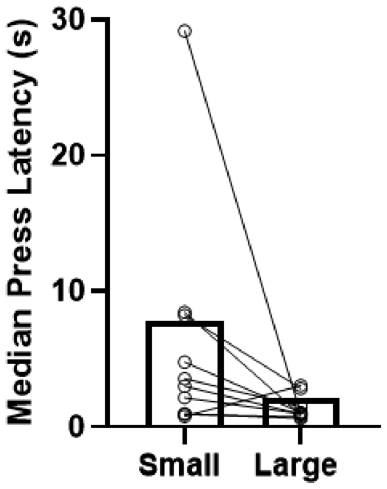


**Figure S2. Analysis of median press latency for Large and Small choices in the absence of delays.** Median press latency on the last day of No-Delay training was significantly shorter for the Large vs Small lever. Wilcoxon test, $p = 0.0273$. $N = 10$ mice.

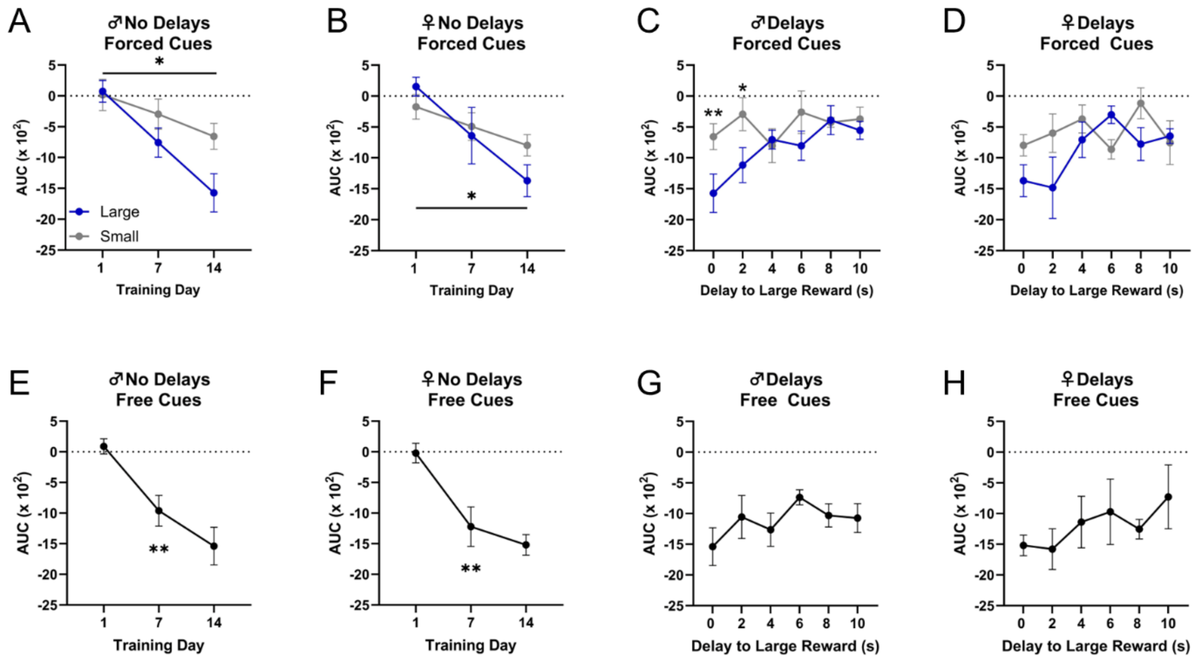


**Figure S3. Analysis of cue-evoked ACh dips by sex in No-Delay training and delay discounting phases. A, B** Both males and females showed a training-dependent increase in ACh dip magnitude (Males: day effect $F_{(2, 20)} = 12.36$, p=0.0003; reward size $F_{(1, 10)} = 4.302$, p = 0.0648; day x reward size $F_{(2, 20)} = 2.186$, p = 0.1385; Females: day $F_{(2, 12)} = 12.73$, p=0.0011; reward size $F_{(1, 6)} = 0.2200$, p=0.6556; day x reward size $F_{(2, 12)} = 2.245$, p=0.1485). Tukey's post hoc: showed differences between sessions 1 and 14 for the large reward for both sexes (Males,*p = 0.0002; females, *p = 0.0007). **C** In males, the mean ACh dip AUC evoked by Large and Small forced cues differed significantly depending on the delay to the large reward. Two-way RM ANOVA: delay $F_{(5, 50)} = 3.476$, p=0.0090; reward size $F_{(1, 10)} = 3.053$, p=0.1112; delay x reward size $F_{(5, 50)} = 2.460$, p=0.0455. Tukey's post hoc: Large vs. Small at 0-s delay (*p = 0.0177) and at 2-s delay (*p = 0.0177). **D** In females, the delay effect and delay x reward size interaction approached statistical significance: delay $F_{(2.903, 17.42)} = 2.878$, p=0.0671; reward size $F_{(1, 6)} = 1.366$, p=0.2869, p=0.1112; delay x reward size $F_{(2.903, 17.42)} = 2.882$, p=0.0669. **E, F** Both males and females show a significant training day effect on the magnitude of the ACh dip evoked by the dual cues in free choice trials in the No-Delay phase (Males: $F_{(2, 10)} = 13.06$, **p=0.0016; females: $F_{(2, 6)} = 16.75$, **p=0.0035). **G, H.** In the delay phase, no significant effects of delay on AUC were found for either sex (Males: $F_{(5, 25)} = 1.583$, p = 0.2012; females: $F_{(5, 15)} = 1.010$, p = 0.4456). Data shown as mean ± SEM. For all panels, n = 6 males and n = 4 females.

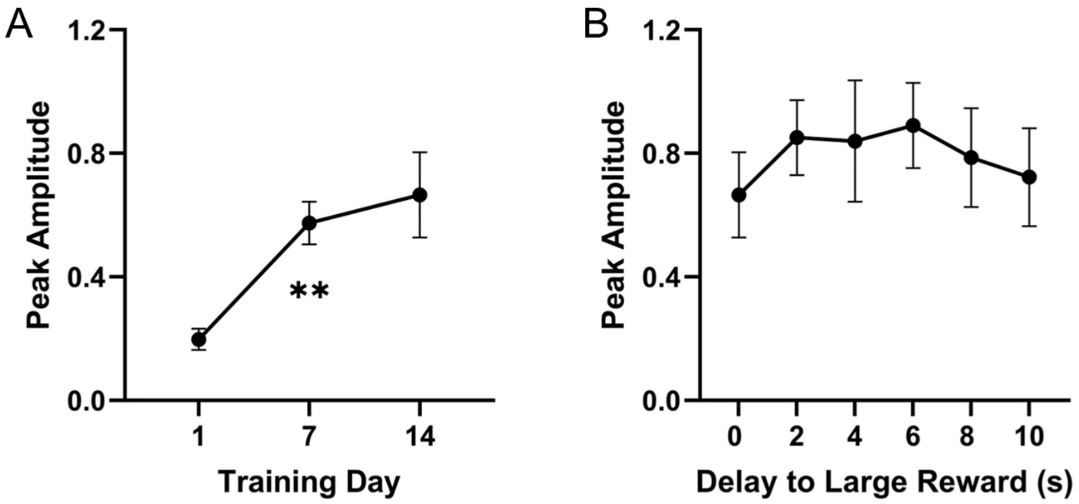


**Figure S4. Analysis of cue-evoked ACh peaks during free choice trials. A** During training, presentation of both cues during the free choice portion of the task significantly increased ACh peak amplitude measured from 0-1s following cue onset: $F_{(2, 18)} = 9.430$, $^{**}p = 0.0016$. **B** Delays did not alter the ACh peak from 0-1s during free choice presentation of both cue lights $F_{(5, 45)} = 0.6877$, $p = 0.6353$. Data shown as mean ± SEM. N = 10 mice.

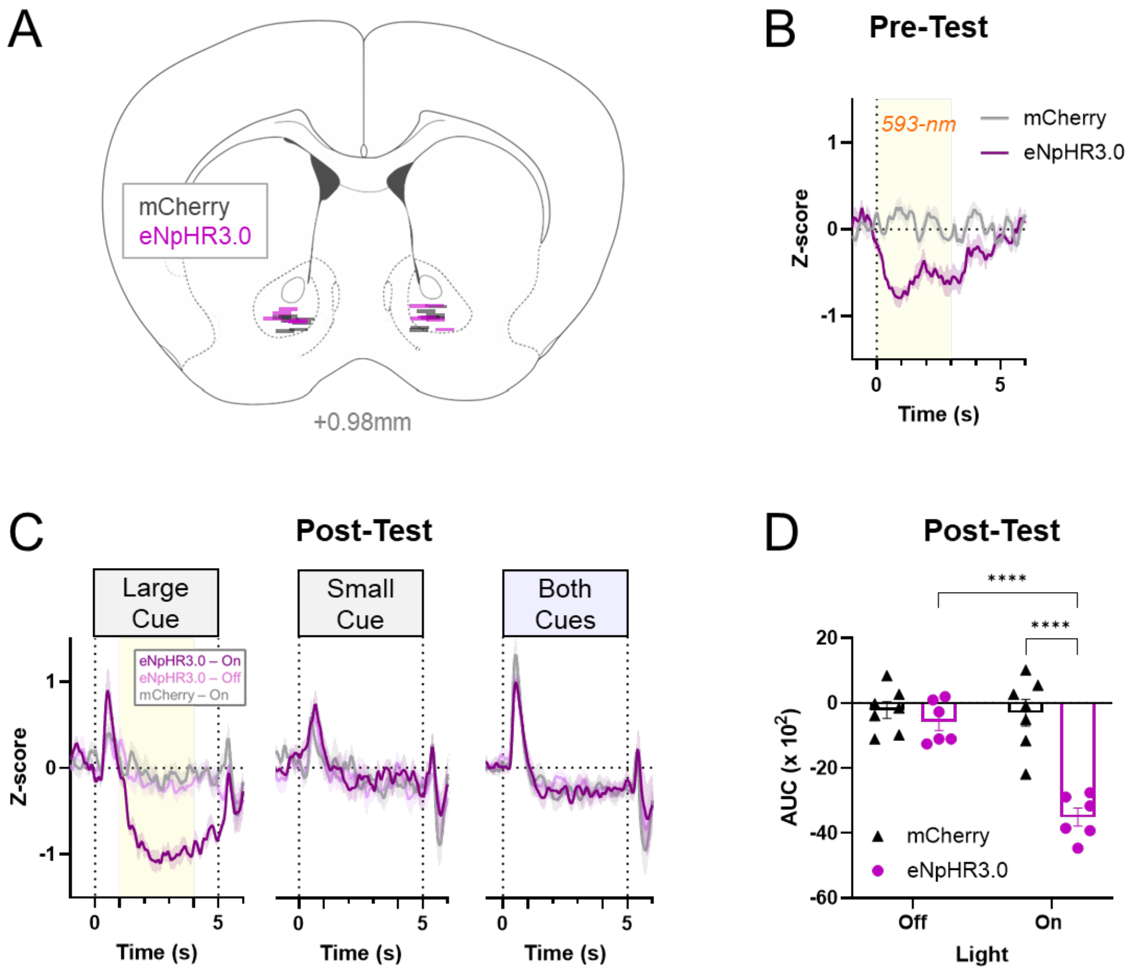


**Figure S5. Validation of optogenetic control of NAc ACh. A** Approximate bilateral fiber locations for eNpHR3.0 and mCherry mice used in optogenetics experiments. **B** Prior to any behavioral training (pre-test), ACh signals were recorded in response to 10 trials of a constant 3-s 593-nm light pulse with a ramped-down shutoff (*see Methods*). Shown are the mean ACh z-scored signals across bilateral responses from all mice showing sustained ACh reduction during the pulse that returns to baseline without a rebound effect. **C** Cue-evoked ACh responses averaged across both hemispheres during a 10-s delay discounting session performed after experiment completion (post-test). *Left*, Large forced cue-evoked ACh responses. 593-nm light illumination was applied 1 second after the cue onset and held constant for 3 s, as described above. Optoinhibition effectively enlarged the cue-evoked ACh dip in eNpHR3.0 but not mCherry mice. Forced Small (*Center*) or free choice *(Right)* cue-evoked ACh responses were not altered by optoinhibition triggered in Large forced trials. Cue-evoked ACh peaks were not affected. **D** Quantification of Large cue-evoked ACh dip in eNpHR3.0 mice relative to control mCherry mice across Light-On and Light-Off sessions. Two-way RM ANOVA revealed main effects of light condition ($F_{(1, 11)}$ = 33.64, p=0.0001) and virus ($F_{(1, 11)}$ = 23.55, p=0.0005), and a significant Light condition x virus interaction ($F_{(1, 11)}$ = 30.12, p = 0.0002). *Post hoc* Fisher's LSD test: mCherry vs. eNpHR3.0 in the Light-On condition and between eNpHR3.0 Light Off vs. On (****p = <0.0001). n = 7 (mCherry); 6 (eNpHR3.0).
